# BAF facilitates interphase nuclear membrane repair through recruitment of nuclear transmembrane proteins

**DOI:** 10.1101/2020.01.10.902296

**Authors:** Alexandra M. Young, Amanda L. Gunn, Emily M. Hatch

## Abstract

Nuclear membrane rupture during interphase occurs in a variety of cell contexts, both healthy and pathological. Membrane ruptures can be rapidly repaired, but these mechanisms are still unclear. Here we show BAF, a nuclear envelope protein that shapes chromatin and recruits membrane proteins in mitosis, also facilitates nuclear membrane repair in interphase, in part through recruitment of the nuclear membrane proteins emerin and LEMD2 to rupture sites. Characterization of GFP-BAF accumulation at nuclear membrane rupture sites confirmed BAF is a fast, accurate, and persistent mark of nucleus rupture whose kinetics are partially dictated by membrane resealing. BAF depletion significantly delayed nuclear membrane repair, with a larger effect on longer ruptures. This phenotype could be rescued by GFP-BAF, but not by a BAF mutant lacking the LEM-protein binding domain. Depletion of the BAF interactors LEMD2 or emerin, and to a lesser extent lamin A/C, increased the duration of nucleus ruptures, consistent with LEM-protein binding being a key function of BAF during membrane repair. Overall our results suggest a model where BAF is critical for timely repair of large ruptures in the nuclear membrane, potentially by facilitating membrane attachment to the rupture site.

## Introduction

The nuclear envelope (NE) is a dynamic protein and membrane compartment that can lose and regain compartmentalization during interphase. Interphase nuclear membrane rupture generally occurs when mechanical stress causes chromatin or nucleoplasm to herniate through gaps in the nuclear lamina, the intermediate filament meshwork that provides mechanical support to the membrane. Tension on the unsupported membrane then leads to membrane rupture (Houthaeve *et al.*, 2018). Nuclear membrane rupture has been observed in a wide range of systems, including cells expressing laminopathy mutations in culture and *in vivo* (De Vos *et al.*, 2011; Earle *et al.*, 2019), cancer cells (Vargas *et al.*, 2012; Yang *et al.*, 2017), cells under mechanical stress in culture and in vivo (Tamiello *et al.*, 2013; Maciejowski *et al.*, 2015; Denais *et al.*, 2016; Hatch and Hetzer, 2016; Raab *et al.*, 2016; Stephens *et al.*, 2017; Xia *et al.*, 2018), and during early *C. elegans* development (Penfield *et al.*, 2018), and ruptures are routinely repaired and generally non-lethal. However, recent studies suggest loss of nuclear compartmentalization can alter transcription (De Vos *et al.*, 2011), and cause mislocalization of large organelles (De Vos *et al.*, 2011; Vargas *et al.*, 2012) and DNA damage (Maciejowski *et al.*, 2015; Denais *et al.*, 2016; Raab *et al.*, 2016; Irianto *et al.*, 2017; Takaki *et al.*, 2017; Pfeifer *et al.*, 2018; Stephens *et al.*, 2019). Based on these data, nuclear membrane rupture is emerging as a major mechanism of genome instability and cell death in lamin-associated diseases and cancer (Isermann and Lammerding, 2017; Houthaeve *et al.*, 2018).

Efficient repair of the nuclear membrane is likely critical for cell viability after rupture and many proteins active in post-mitotic NE assembly localize to nucleus rupture sites (reviewed in (LaJoie and Ullman, 2017). One such protein is BAF (barrier-to-autointegration factor), which accumulates at nucleus rupture sites (Denais *et al.*, 2016; Raab *et al.*, 2016; Penfield *et al.*, 2018; Halfmann *et al.*, 2019) and on mitotic chromatin, where it cross-links chromatin into a single mass (Samwer *et al.*, 2017), and recruits LEM-domain (Lap2, emerin, Man1-domain) NE transmembrane proteins (NETs) and lamin A (Jamin and Wiebe, 2015). A recent paper suggested BAF is also essential for interphase nuclear membrane repair via recruitment of membrane bound proteins to rupture sites (Halfmann *et al.*, 2019). However, it remains unknown how broad the requirement for BAF is, and whether it is similarly required to repair ruptures that occur in cells where nuclear lamina organization is perturbed, as in the presence of laminopathy and cancer mutations (De Vos *et al.*, 2011; Yang *et al.*, 2017; Earle *et al.*, 2019; Zhang *et al.*, 2019).

To address these questions, we turned to a nuclear membrane rupture system based on depletion of LMNB1 in U2OS cells. Cell cycle arrest causes individual nuclei to undergo multiple spontaneous ruptures of variable extent and our previous work suggested this cell line recapitulates all the major features of nucleus rupture and repair (Hatch and Hetzer, 2016). Our current data demonstrate that although BAF is a sensitive, reliable, and long-lasting marker of membrane rupture sites, it is not required for membrane repair. Instead we observe a substantial lengthening of rupture duration after BAF depletion, with longer ruptures being more likely to be affected. Additional experiments with a LEM-domain binding mutant of BAF and depletion of the BAF interacting proteins emerin, LEMD2, and lamin A/C, indicate BAF functions by recruiting LEM-domain proteins, which is likely especially critical to repair large nuclear membrane gaps.

## Results and Discussion

To characterize BAF kinetics in our system we stably expressed GFP-BAF, 2xRFP-NLS (RFP-NLS (nuclear localization signal)) and shRNAs against LMNB1 (shLMNB1) in U2OS cells. Depletion of lamin B1 (Figure S1A) caused an increase in the number and size of nuclear lamina gaps, as expected (Figure S1B), and did not affect rupture and repair kinetics (Figure S1C). These kinetics were analyzed by comparing RFP-NLS intensity changes over time. RFP-NLS is nuclear when the membrane is intact, upon rupture it is visible in the cytoplasm as a diffuse signal that is initially brightest proximal to the rupture site, and it is reimported into the nucleus as nucleus integrity is restored (Movie S1). Nucleus rupture is frequently accompanied by a quantifiable decrease in nuclear RFP-NLS intensity and the proportion decrease is defined as the rupture extent (Denais *et al.*, 2016; Raab *et al.*, 2016; Deviri *et al.*, 2019; Zhang *et al.*, 2019). To compare repair kinetics between control and LMNB1 depleted cells, analysis was limited to ruptures of similar extents in both populations to control for the duration of RFP-NLS reimport.

GFP-BAF kinetics during rupture and repair were assessed in U2OS RFP-NLS shLMNB1 GFP-BAF cells arrested in S phase for 24 hr to increase the frequency of membrane ruptures (Hatch and Hetzer, 2016). Cells were imaged either every 30 sec or every 3 min and total RFP-NLS and GFP-BAF rupture site intensities were analyzed over time. We found GFP-BAF was rapidly (within 30 sec) recruited to membrane rupture sites, followed by a gradual decrease in intensity until a plateau was reached (Figure 1, A and B; Movie S1). This plateau frequently remained higher than GFP-BAF intensity in the surrounding NE (50/58 ruptures) for several hours after RFP-NLS re-accumulation (Figure 1A). We observed distinct GFP-BAF foci for each rupture, even when multiple ruptures clustered spatially or temporally (Figure S1, D and E), or the rupture extent was too small to quantify (Figure S1E). Within an individual nucleus, the peak intensity of GFP-BAF frequently correlated with rupture extent (12/15 cells, Figure 1B), as previously observed (Denais *et al.*, 2016). Consistent with studies in different systems (Denais *et al.*, 2016; Halfmann *et al.*, 2019), our results demonstrate GFP-BAF is a sensitive and accurate marker of nucleus rupture that marks the site of cytoplasmic-exposed chromatin for a substantial time after membrane repair. In addition, similar to lamin A “scars” (Denais *et al.*, 2016), rupture-induced BAF foci appear refractory to re-rupture.

**Figure 1.**
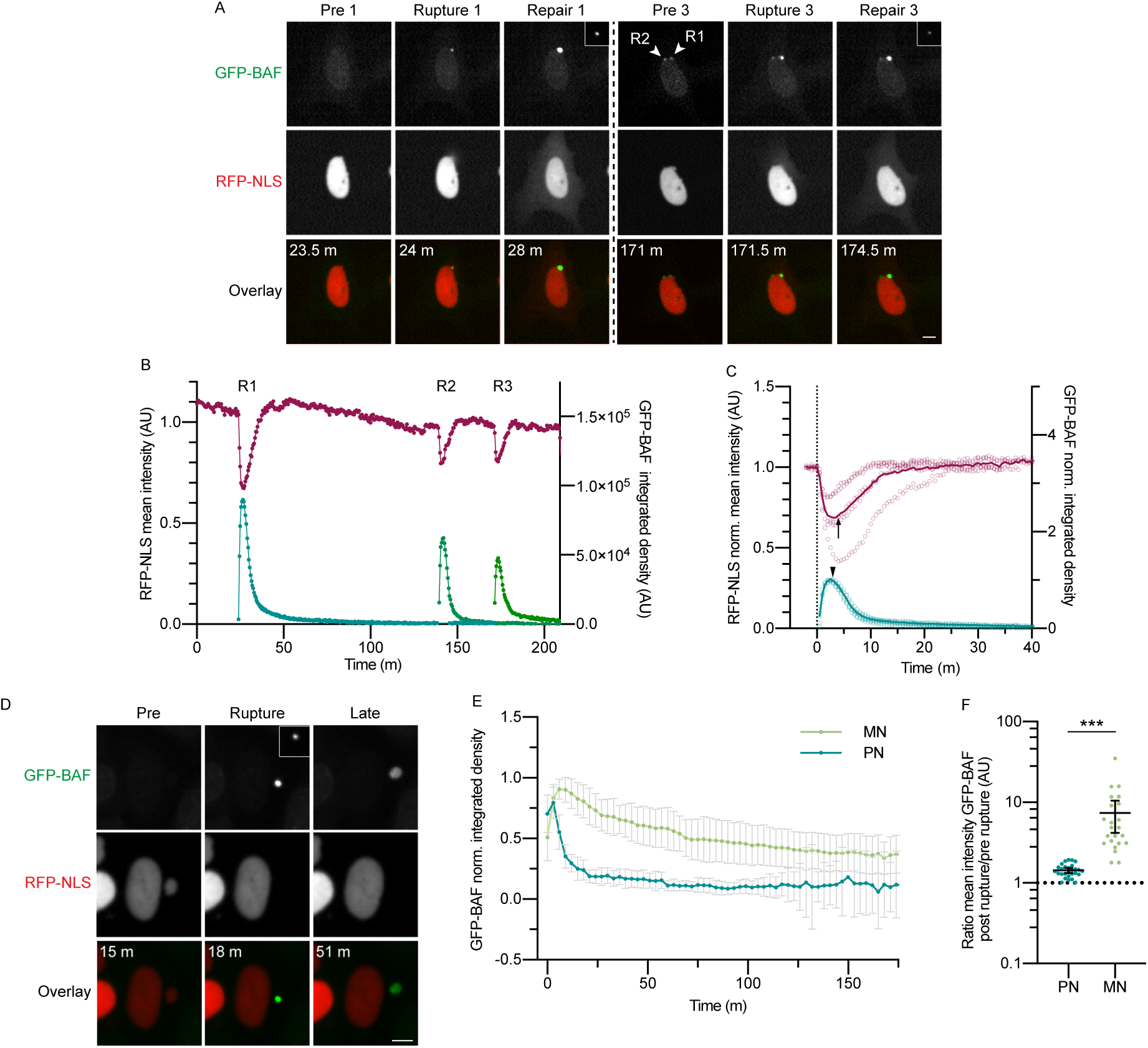
Characterization of GFP-BAF accumulation and loss kinetics at membrane rupture sites. (A) Still images of U2OS GFP-BAF RFP-NLS shLMNB1 cells imaged every 30 sec. RFP-NLS images gamma adjusted to increase visibility of cytoplasmic RFP, insets are unsaturated images of GFP-BAF at rupture sites (same size). m = minutes post imaging start. Two of three ruptures are depicted, GFP-BAF foci from previous ruptures (R1, R2) are marked with arrowheads in image prior to R3. Rupture = first frame with cytoplasmic RFP-NLS, repair = first frame of RFP-NLS cytoplasmic intensity decrease. (B) RFP-NLS mean intensity (magenta) and GFP-BAF integrated density (green) traces of ruptures in (A). GFP-BAF quantified only at rupture-associated foci. (C) RFP-NLS and GFP-BAF intensity traces from 5 ruptures between 2.5 and 5 min duration, 30 sec pass time. GFP-BAF values normalized to max intensity. Mean (thick line) and replicates (open circles) are plotted. Arrow = time of repair start, arrowhead = time of first GFP-BAF decrease. (D) Still images of micronucleus rupture. Inset is unsaturated image of GFP-BAF (same size). (E) GFP-BAF intensity traces from micronuclei (MN) or primary nuclei (PN) ruptures. Traces were normalized to the GFP-BAF max intensity prior to averaging. Average (mean) intensity and 95% CI are plotted. N values: MN, 8; PN 13 ruptures. (F) Ratio of GFP-BAF mean intensity at post rupture (and repair -PN) plateau over mean intensity pre rupture (5 frame average) (n values: PN, 28; MN, 23). ***p<0.001, K-S test. Median and 95% CI are plotted. Both populations were significantly different from 1, p<0.001, One-sample t-test. All scale bars = 10 μm. AU = arbitrary units.

Analysis of the kinetics of GFP-BAF recruitment demonstrated GFP-BAF levels started to decline prior to the start of RFP-NLS regain in 24/24 ruptures (Figure 1C and Figure S1F, arrowheads, only ruptures >3 min included), consistent with a previous report (Denais *et al.*, 2016). To define the effect of membrane repair on BAF kinetics, we analyzed GFP-BAF recruitment to ruptured micronuclei. Micronuclei (MN) form in human cells when chromosomes missegregate during mitosis and recruit their own NE. In contrast to the “primary” nucleus, nuclear membrane ruptures in MN are rarely able to repair (Hatch *et al.*, 2013). Analysis of live-cell imaging of micronucleated cells, generated by addition of the spindle checkpoint inhibitor reversine (Santaguida *et al.*, 2010), showed GFP-BAF accumulated on ruptured MN concurrently with the loss of RFP-NLS and remained accumulated on the exposed chromatin until the end of imaging or mitosis (Figure S1G, Movie S1) (Liu *et al.*, 2018). In addition, GFP-BAF frequently accumulated at a single focus on the MN and spread throughout the chromatin (Figure 1D, 53/56 MN). GFP-BAF loss was significantly slower and the end intensity higher on ruptured MN compared to repairing primary nuclei (Figure 1, E and F, Movie S1). Together these data suggest GFP-BAF removal from rupture sites is initiated independently of nucleus re-compartmentalization, but further loss is driven by membrane repair. Our data are consistent with binding of non-phosphorylated cytoplasmic BAF to the exposed chromatin and then release by inhibitory phosphorylation of BAF by nuclear kinases (Halfmann *et al.*, 2019). The increased BAF release after nucleus integrity regain could reflect a general increase in kinase concentrations and/or targeted removal of BAF by the reforming membrane, similar to what may occur during NE assembly (Samwer *et al.*, 2017).

We next examined the function of BAF in nuclear membrane rupture and repair. U2OS RFP-NLS shLMNB1 cells were transfected with siRNAs against BAF (siBAF) or a control siRNA (siControl) for 24 hr, arrested in S phase for 24 hr, then imaged every 3 min. BAF protein levels were strongly reduced 48 hr after siRNA transfection (Figure 2B), but BAF-associated mitotic phenotypes, including extensive mislocalization of lamin A/C and emerin and multinucleation (Haraguchi *et al.*, 2008; Samwer *et al.*, 2017), were absent (Figure S2, A and B). Analysis of nucleus rupture frequency showed no difference between control and BAF depleted cells (Figure S2, C and D). Surprisingly, BAF depletion also did not affect the ability of nuclei to re-compartmentalize; 98% (84/86) of siBAF cells, versus 97% (99/102) of siControl cells, fully recovered RFP-NLS after each rupture. In contrast, analysis of the duration of individual ruptures, defined as the time RFP-NLS is visible in the cytoplasm (Figure 2A), demonstrated BAF depletion significantly increased the duration of nucleus rupture compared to control cells in both shLMNB1 (Figure 2C, Movie S1) and normal U2OS cells (Figure S2, E and F). These data demonstrate BAF is not required for nucleus membrane repair but does significantly increase its efficiency.

**Figure 2.**
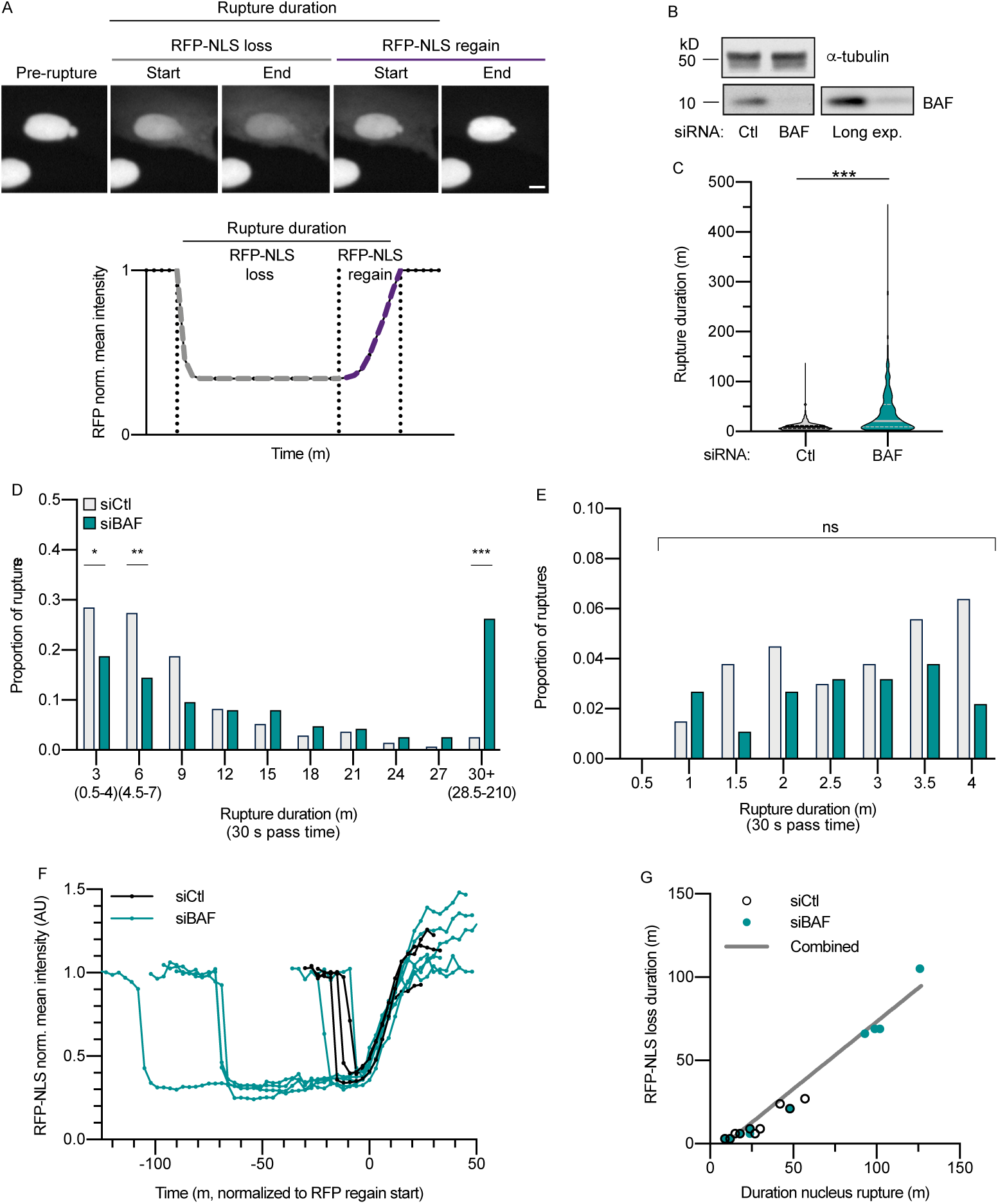
BAF depletion delays nuclear membrane repair. (A) Top: Still images of U2OS RFP-NLS shLMNB1 cells. RFP-NLS is gamma adjusted. Scale bar = 10 μm. Bottom: Cartoon of normalized RFP-NLS mean intensity trace during a nucleus rupture. Rupture duration is defined from images by multiplying the number of images with cytoplasmic RFP-NLS by 3 min and RFP-NLS intensity loss and regain are defined by the trace as indicated and described in the text. (B) Representative immunoblot of BAF protein levels 48 hr after siRNA transfection. A longer exposure is shown at right. (C) Quantification of nucleus rupture duration in cells treated with the indicated siRNAs (n values: siCtl, 416; siBAF, 387 ruptures over 3 experiments). ***p<0.001, K-S test. (D) Histogram of proportion of ruptures with indicated nucleus rupture durations from cells transfected with indicated siRNAs, imaged every 30 sec, binned to every 3 minutes (representative bin durations indicated below) (n values: siCtl, 266; siBAF 186, ruptures over 3 experiments). ***p<0.001, **p<0.01, *p<0.05, Fisher’s exact test. Statistics for ruptures between 9 and 30 min are omitted for readability. (E) Expansion of nucleus rupture duration values binned into “3 min” in (D). ns = p>0.05, χ^2^ test. (F) RFP-NLS trace analysis of large extent ruptures (>0.6) from cells transfected with siCtl or siBAF. t_0_ = RFP-NLS regain start (A) (n values: siCtl, 3; siBAF, 6 ruptures) (G) RFP-NLS intensity loss duration compared to nucleus rupture duration from siCtl and siBAF cells (n values: siCtl, 10; siBAF, 9 ruptures). Best fit line for the combined data sets is shown.

To determine whether the result was specific to this nucleus rupture mechanism, we assessed the effect of BAF depletion on membrane repair after laser-induced nucleus rupture. Unexpectedly, we found BAF depletion did not increase rupture duration in this context (Figure S2, G and H). However, the laser-induced ruptures had a significantly smaller extent yet took much longer to re-compartmentalize than spontaneous ones in control cells (Figure S2, I-K). Based on these altered rupture characteristics, and the likelihood that laser wounding altered protein structures at rupture sites, we decided to focus only on spontaneous ruptures.

We next compared the proportion of ruptures at each duration between BAF depleted and control cells to determine whether all ruptures are equally dependent on BAF for repair. To facilitate analysis, we binned together ruptures lasting 30 min or longer (top 5% of control ruptures). We expected if BAF depletion affects all ruptures equally, we would see a translation of the histogram towards longer durations. However, an initial analysis showed BAF depletion only modestly altered the proportion of 3 min ruptures compared to the next longest ones (6 min) and ruptures lasting 30 min or over (Figure S2L). The 3 min ruptures in BAF-depleted cells frequently occurred in the same nuclei as 30 min+ ruptures (Figure S2M), indicating this result is unlikely due to transfection efficiency. To obtain a higher temporal resolution of short duration ruptures, since those captured by a single frame could have a duration between 200 msec to nearly 6 min, we imaged siControl and siBAF transfected cells every 30 sec. When rupture durations were partitioned into 3 min bins, we observed the same trends as with a 3 min imaging pass time, indicating our standard conditions capture the majority of short ruptures (Figure 2D). Comparing the relative frequency of ruptures between 0.5 and 4 min duration showed no significant difference between the BAF depleted and control cell ruptures (Figure 2E), suggesting rapidly repairing membrane ruptures are less dependent on BAF compared to longer ones.

Delay in RFP-NLS regain after BAF depletion could be due to defects in initiating membrane repair (or other mechanisms of re-compartmentalization), defects in membrane sealing, or altered nuclear import. To distinguish between these hypotheses, we analyzed the timing and kinetics of RFP-NLS intensity loss and regain in siBAF and siControl transfected cells (Figure 2A). If initiation of membrane repair were defective, then we expected the duration of RFP-NLS intensity loss to lengthen in response to BAF depletion, whereas if membrane resealing or nuclear import were altered, the duration of RFP-NLS intensity regain should lengthen. Analysis of RFP-NLS regain duration found that it generally correlated with rupture extent in both populations (Figure S2N), confirming the necessity of comparing ruptures of similar extents to evaluate repair kinetics. Overlaying RFP-NLS intensity traces from siControl and siBAF ruptures of similar extent found the duration of RFP-NLS loss, not the duration of RFP-NLS regain, increased with increasing rupture duration (Figure 2F). Analyzing RFP-NLS loss duration over a wide range of rupture durations showed this correlation was independent of BAF depletion (Figure 2G). Together, these data strongly suggest loss of BAF delays the initiation of re-compartmentalization during ruptures lasting longer than a few minutes.

To determine whether BAF interactions with LEM-domain NETs are required for its activity in membrane repair, we first evaluated the ability of a mutated BAF that lacks the LEM-domain binding site (L58R (Samwer *et al.*, 2017)) to rescue BAF depletion compared to wild-type BAF. Stably expressed WT-GFP-BAF and L58R-GFP-BAF were resistant to siRNAs against endogenous BAF (Figure 3A), and localized to the NE and membrane rupture sites in interphase (Figure 3, B and C), with L58R-GFP-BAF showing decreased recruitment to the NE compared to WT, as expected (Samwer *et al.*, 2017). WT-GFP-BAF rescued nucleus rupture duration after BAF depletion to control levels (Figure 3D), demonstrating that the BAF siRNA was specific. In contrast, L58R-GFP-BAF expression failed to rescue rupture duration compared to control cells (Figure 3D), and had only a slightly lower proportion of 30 min+ ruptures than siBAF plus GFP alone cells (Figure 3E). These data suggest the LEM-binding region of BAF is a critical component of its membrane repair function.

**Figure 3.**
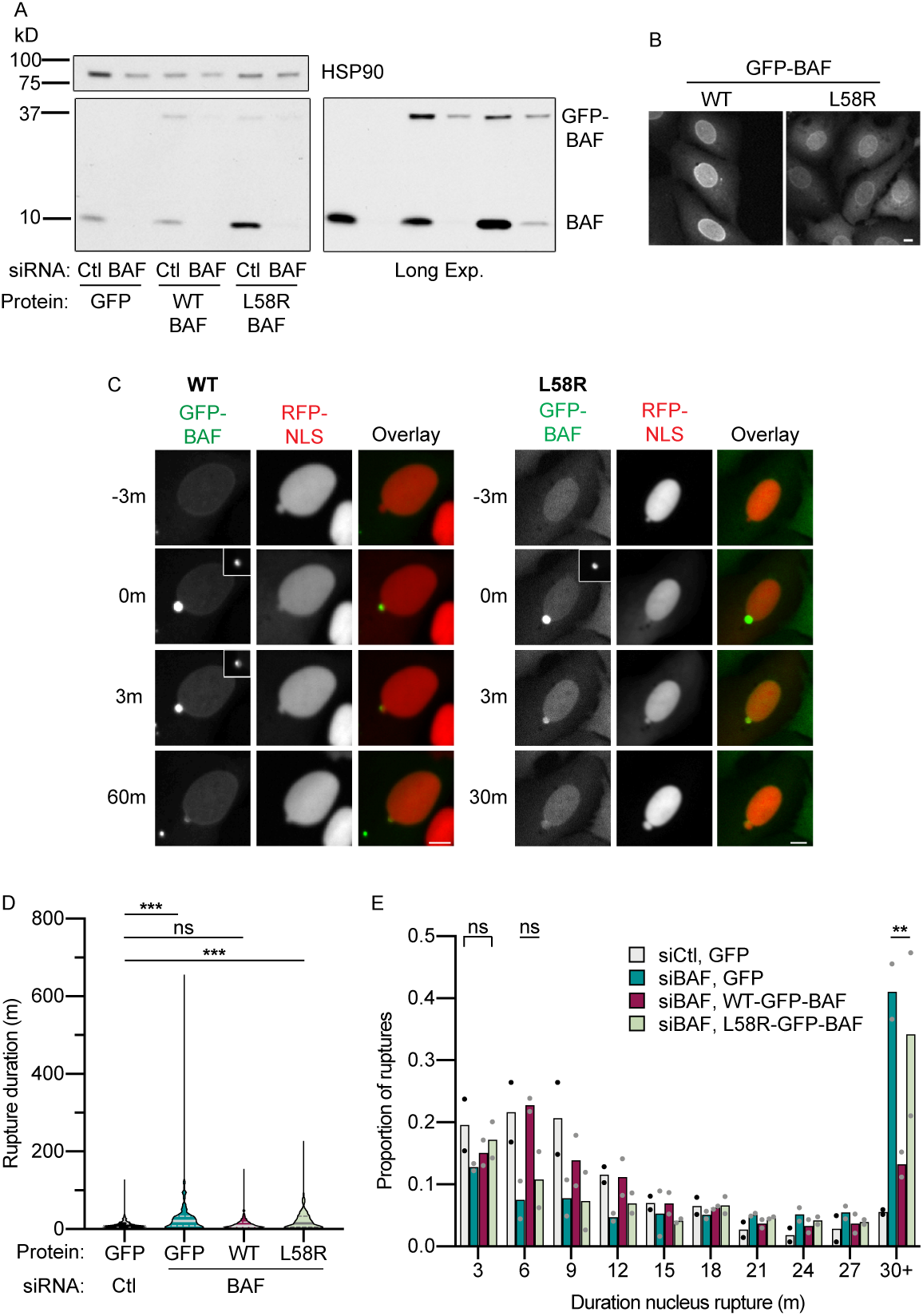
BAF LEM-protein binding function is required for efficient nuclear membrane repair. (A) Representative immunoblot of BAF protein levels in U2OS RFP-NLS shLMNB1 expressing indicated proteins (all GFP-BAF) 48 hr after depletion of endogenous BAF. A longer exposure (Long Exp.) is shown at right. (B) Representative images of non-transfected cells expressing either WT- or L58R-GFP-BAF. (C) Still images of cells expressing either WT- or L58R-GFP-BAF 48 hr after transfection with siBAF. T_0_ = rupture start. Insets are unsaturated images of GFP-BAF at rupture sites. Scale bars = 10 μm. (D) Quantification of nucleus rupture durations in cells expressing indicated proteins and transfected with siBAF. N values: siCtl+GFP, 237; siBAF+GFP, 235; siBAF+GFP-BAF, 228 siBAF+L58R-GFP-BAF, 321 ruptures from 2 experiments. ***p<.001, ns=p>0.05, K-W test. (E) Histogram of proportion of ruptures shown in (D) with indicated rupture durations. Bracket includes all conditions, lines indicate pairwise comparison between siBAF, GFP and siBAF, L58R-GFP-BAF cells. ns = p>0.05, χ^2^ test (bracket) or Fisher’s exact test (line). **p<0.01, Fisher’s exact test.

BAF interacts with several proteins that accumulate at rupture sites, including lamin A/C, emerin, and LEMD2 (Denais *et al.*, 2016; Halfmann *et al.*, 2019). To determine which BAF binding partner(s) contributes to efficient membrane repair, we first evaluated whether these proteins require BAF to accumulate at membrane ruptures. Using cytoplasmic RFP-NLS to identify ruptured nuclei, we found severely reduced recruitment of lamin A/C, emerin, and LEMD2 to rupture sites in BAF depleted cells (Figure 4, A and B), confirming previous results (Halfmann *et al.*, 2019). Live-cell imaging of BAF depleted cells expressing mCherry-lamin A showed lamin A accumulation at rupture sites was absent, not delayed (Figure S3, A and B). Although it is likely emerin and LEMD2 accumulation is also inhibited, we cannot rule out delayed recruitment in the absence of BAF.

**Figure 4.**
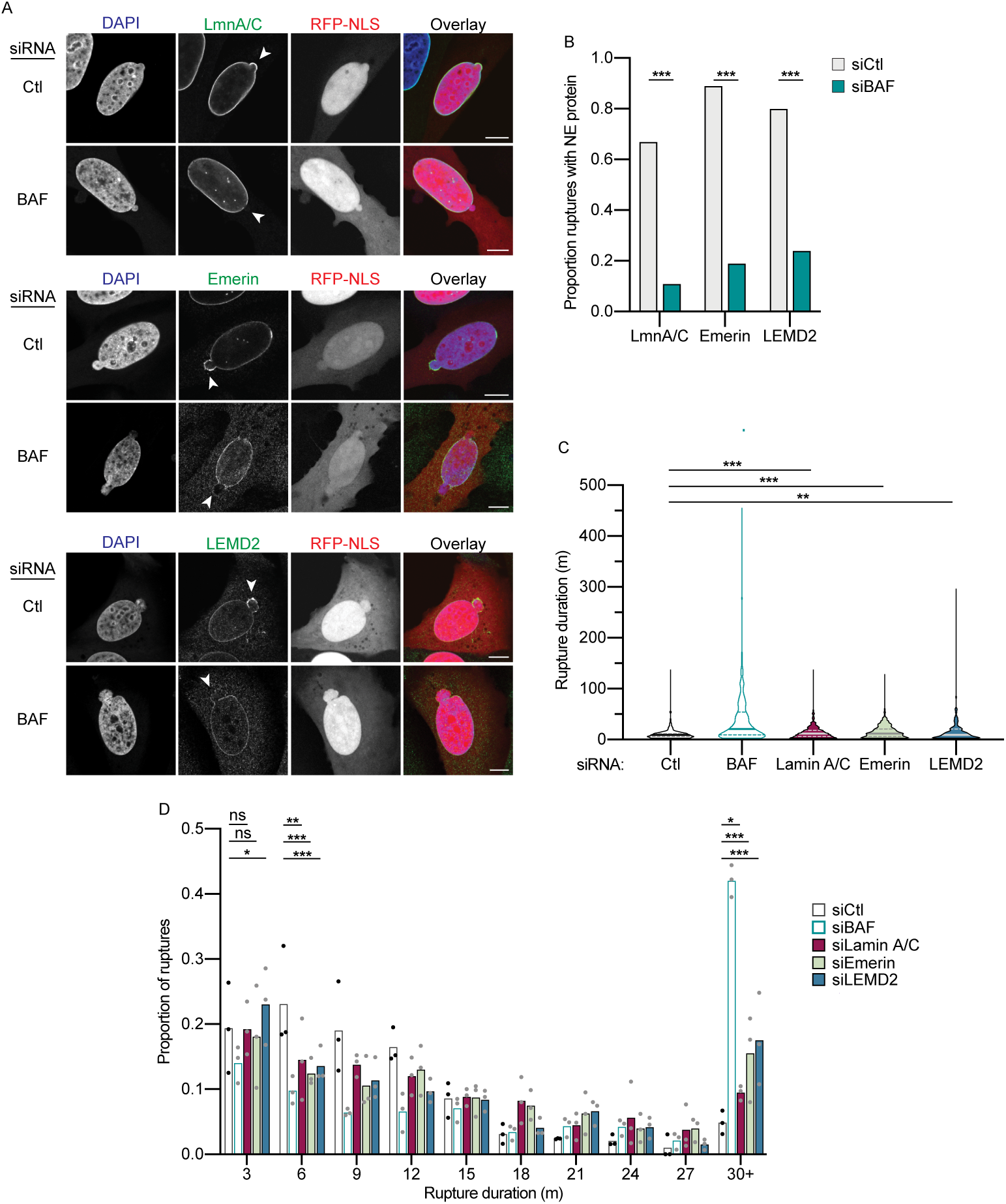
BAF recruits lamin A/C, emerin and LEMD2 to promote nucleus re-compartmentalization. (A) Representative fixed images of U2OS RFP-NLS shLMNB1 cells transfected with siControl (siCtl) or siBAF undergoing nuclear membrane rupture, determined by cytoplasmic RFP-NLS. Cells were labeled with antibodies to lamin A/C, emerin, or LEMD2. RFP-NLS images are gamma adjusted. Scale bars = 10 μm. Proportion of rupture sites with accumulated lamin A/C (n values: siCtl, 50; siBAF, 55), emerin (siCtl, 62; siBAF, 70), or LEMD2 (siCtl, 92; siBAF, 95). Arrowheads = estimated rupture site. N-values are pooled from 3 experiments. ***p<0.001, Fisher’s exact test. (C) Quantification of nucleus rupture durations after transfection with siRNAs against LMNA, EMD, or LEMD2. Data for siCtl and siBAF collected at the same time are reproduced from figure 2C (open bars) (n values: siLMNA, 548; siEmerin, 416; siLEMD2, 887 ruptures from 3 experiments). ***p<0.0001, K-W test. (D) Histogram of proportion of ruptures shown in (C) with indicated durations of nucleus rupture. ***p<0.001, **p<0.01, *p<0.05, ns p>0.05, Fisher’s exact test.

To determine whether lamin A/C, emerin, or LEMD2 function in nuclear membrane repair, U2OS RFP-NLS shLMNB1 cells were transfected with siRNAs against each protein individually and nucleus rupture duration analyzed in S-phase arrested cells by live-cell imaging 48 hr after the initial siRNA transfection. siRNA transfection was sufficient to deplete the targeted proteins to at least 50% of control levels by western blot (Figure S3C) and most cells showed little to no protein expression by immunofluorescence (Figure S3, D-F).

Depletion of any one of these proteins caused a statistically significant increase in the median nucleus rupture duration (Figure 4C), with depletion of emerin or LEMD2 also causing a substantial increase in the proportion of ruptures longer than 30 min (Figure 4D). Consistent with our BAF depletion results, loss of lamin A/C, emerin, or LEMD2 showed less decrease in the proportion of 3 min ruptures compared to 6 min ruptures, with LEMD2 depletion instead causing a significant increase in 3 min ruptures (Figure 4D). Thus, our results suggest BAF-dependent recruitment of NE proteins to membrane rupture sites is required for efficient repair of longer ruptures.

Our data suggest BAF is critical for membrane repair for a large proportion of interphase nucleus ruptures driven by nuclear lamina disorganization and nucleus compression, and that this activity is due to BAF-dependent accumulation of NETs at rupture sites. Although knockdown of any one of three BAF-interacting proteins, lamin A/C, emerin, or LEMD2, caused a significant increase in rupture duration, only emerin and LEMD2 depletion caused a substantial increase in the proportion of very long, 30 min+, ruptures. In addition, a small increase in rupture duration after lamin A/C depletion is consistent with previous reports that lamin A/C depletion increases rupture extent (Denais *et al.*, 2016; Zhang *et al.*, 2019). Thus, in spite of discrepancies between our observations, both our results and those in Halfmann *et al.*, (2019) support a general model where BAF promotes nucleus re-compartmentalization by binding NETs and new membrane to the exposed chromatin. Additionally, our observation that BAF depletion does not eliminate very short ruptures (3 min or less) suggests different types of ruptures have different requirements for re-compartmentalization. >We propose the main difference is the size of the membrane gap and larger gaps have a higher requirement for BAF-dependent protein and membrane recruitment. Determining whether a similar mechanism, such as direct interactions between NETs and chromatin (Barton *et al.*, 2015), repairs both small ruptures and large ruptures in the absence of BAF, or whether small rupture membrane repair occurs by a currently uncharacterized mechanism will be an important question for future studies that will provide new insight into how nuclear membrane dynamics are regulated.

## MATERIALS AND METHODS

### Cell culture and construction of stable cell lines

U2OS cells were cultured in Dulbecco’s modified Eagle medium (DMEM; GIBCO) supplemented with 10% (v/v) fetal bovine serum (FBS; GIBCO), and 1% (v/v) penicillin-streptomycin (Sigma-Aldrich) at 37°C with 10% CO2 in a humidified incubator. For nucleus rupture experiments, cells were incubated with 2 mM hydroxyurea (EMD Millipore) for 24 hr prior to fixation and throughout imaging. For micronucleus rupture experiments, cells were incubated with 0.5 µM reversine (Fisher Sci) for 4 hr prior to and throughout imaging. U2OS RFP-NLS cells were made by transfection with 2xRFP-NLS, selecting with 0.5 mg/ml G418 (GIBCO), and FACS enrichment for RFP+ cells. U2OS RFP-NLS EGFP-BAF shLMNB1 cells were made by infecting U2OS RFP-NLS cells with lentiviruses from pLKO.1 shRNA-LMNB1.71 and EGFP-BAF-IRES-Blast, selecting with 10 μg/ml blasticidin (InvivoGen) and 2 μg/ml puromycin (Sigma-Aldrich), and FACS enrichment for RFP/GFP double positive cells. U2OS GFP-NLS mCherry-lamin A shLMNB1 cells were made by infecting U2OS GFP-NLS shLMNB1 cells (Hatch *et al.*, 2013) with lentivirus expressing mCherry-Lamin A, blasticidin selection, and FACS enrichment for RFP/GFP double positive cells.

### Plasmids

Plasmids encoding EGFP-BAF and BAF mutants (EGFP-BAF_WT, EGFP-BAF_L58R, EGFP-BAF_G47E and EGFP-BAF_G25E) were a gift from the Gerlich lab (IMBA, Vienna, Austria) and described in (Samwer *et al.*, 2017). EGFP-IRES-Blast was generated by digesting EGFP-BAF with XbaI and BamHI and inserting EGFP PCR’d from the same construct by sequence- and ligation-independent cloning (SLiC). Lamin B1 shRNA was expressed from pLKO.1 shRNA-LMNB1.71 puro (SHCLND-NM_005573, Sigma-Aldrich) containing the sequence: 5′ CCGGGCATGAGAATTGAGAGCCTTTCTCGAGAAAGGCTCTCAATTCTCATGCTTTTT-3′. 2xRFP-NLS (RFP-NLS) was described previously (Hatch *et al.*, 2013) and contains mCHerry fused to TagRFP and a C-terminal NLS (PPKKKRKV) in a pcDNA backbone. 3xGFP-NLS was described previously (Vargas *et al*., 2012) and contains cycle3GFP fused to EGFP, an NLS, and a second EGFP in a pcDNA backbone. pmCherry-Lamin A was made by PCR and ligation of prelamin A from pBABE-puro-GFP-wt-lamin A (a gift from Tom Misteli (Scaffidi and Misteli, 2008), Addgene plasmid #17662; http://n2t.net/addgene:17662; RRID:Addgene_17662) with an N-terminal mCherry fusion behind an EF1α promoter in a pLVX backbone (Takara Bio). The promoter plus gene fusion was then cloned into the pEGFP-BAF backbone by PCR and ligation, replacing both the crippled EF1α promoter and the EGFP-BAF sequence.

### siRNA transfection

All siRNA transfections were performed using siLentFect (Bio-Rad) according to manufacturer’s instructions. Custom siRNAs (Dharmacon) against human BAF (#1. 5′-AGUUUCUGGUGCUAAAGAAtt-3′, #2. 5′-CCCUCACUUUCAAUCCGUUuu-3′), lamin A/C (5′-GGUGGUGACGAUCUGGGCUuu-3′)(Harada *et al.*, 2014), emerin (5′-GGUGCAUGAUGACGAUCUUtt-3′) (Salpingidou *et al.*, 2007), and LEMD2 (5′-UUGCGGUAGACAUCCCGGG[dT][dT] −3) and a nontargeting control against Luciferase (5′-UAUGCAGUUGCUCUCCAGC[dT][dT] −3′) were used for gene knockdowns. siRNAs were added to 100 nM final concentration. Cells were analyzed 48 hr after a single siRNA transfection at t = 0 hr for all proteins except LEMD2, where cells were transfected at 0 and 24 hr.

### Immunoblotting

Cells were lysed directly in 1x SDS/PAGE sample buffer (LC2570; Thermo Fisher Scientific) + βME and separated on 4–15% gradient gels (Mini-PROTEAN TGX; Bio-Rad) then transferred to nitrocellulose membrane (0.2 μm; Bio-Rad). Membranes were blocked with 5% (w/v) milk and 0.25% (v/v) Tween 20, then incubated with appropriate primary antibodies and HRP or near-IR-conjugated secondary antibodies and visualized using chemiluminescence (SuperSignal West Femto Chemiluminescent Substrate; Thermo Fisher Scientific), or a Li-Cor Odyssey fluorescence scanner, respectively. Only fluorescently imaged blots were quantified using the integrated density and normalizing to loading controls.

#### Primary antibodies (human proteins)

Mouse anti–BAF (1:250; clone A11; sc-166324; Santa Cruz Biotechnology)

Rabbit anti–Lamin A/C (1:10,000; clone EPR4100; ab108595; Abcam)

Mouse anti–Lamin B1 (1:1,000; clone C12; sc-365214; Santa Cruz Biotechnology)

Mouse anti–Emerin (1:500; clone 4G5; EMERIN-CE; Leica Biosystems)

Rabbit anti–LEMD2 (1:1,000; HPA017340; Sigma-Aldrich)

Mouse anti–GAPDH (1:1,000; clone GT239; GTX627408; GeneTex)

Mouse anti–α-Tubulin (1:5,000; clone DM1A, GTX27291; GTX; GeneTex)

Rabbit anti-HSP90 (1:5000; 4874; Cell Signaling Technology)

#### Secondary antibodies

HRP-conjugated goat anti–mouse (1:5,000; G-21040; Thermo Fisher Scientific)

HRP-conjugated goat anti–rabbit (1:5,000; G-21234; Thermo Fisher Scientific)

HRP-conjugated goat anti–rabbit (1:5,000; G-21234; Thermo Fisher Scientific)

Alexa Fluor 790-conjugated donkey anti–rabbit (1:10,000; A-11374; Thermo Fisher Scientific)

Alexa Fluor 790-conjugated donkey anti–mouse (1:10,000; A-11371; Thermo Fisher Scientific)

Alexa Fluor 680-conjugated donkey anti–rabbit (1:10,000; A-10043; Thermo Fisher Scientific)

Alexa Fluor 680-conjugated donkey anti–mouse (1:10,000; A-10038; Thermo Fisher Scientific)

### Immunofluorescence and fixed cell imaging

Cells were grown on poly-L lysine coated coverslips and fixed with freshly made 3% paraformaldehyde (from 16% paraformaldehyde (w/v), EM grade; Electron Microscopy Sciences) in 1x PBS for 10 min at RT. Fixed cells were permeabilized with 1x PBS containing 0.4% Triton X-100 for 15 min, incubated with primary antibody for 1 hr, washed 1x PBS, incubated with secondary antibody for 1 hr, incubated in 0.1 ug/mL DAPI (Life Technologies) for 5 min, and mounted in Vectashield (Vector Laboratories). Imaging was performed using a 40x/1.30 Plan Apo Leica objective on a Leica DMi8 outfitted with a TCS SPE scan head with spectral detection. Images were acquired using the LAS X software platform (version 3.5.5.19976). Images were corrected for brightness and contrast using FIJI (Schindelin *et al.*, 2012) and Photoshop (Adobe). A gamma function was applied to most RFP-NLS images to enhance the visibility of the cytoplasmic signal and noted in the figure legend where used. Images are single sections unless noted in the figure legends.

#### Primary antibodies

Rabbit anti–Lamin A/C (1:1,000; clone EPR4100; ab108595; Abcam)

Mouse anti–Lamin B1 (1:100; clone C12; sc-365214; Santa Cruz Biotechnology)

Mouse anti–Emerin (1:100; clone 4G5; EMERIN-CE; Leica Biosystems)

Rabbit anti–LEMD2 (1:100; HPA017340; Sigma-Aldrich)

Rabbit anti–cGAS (1:100; clone D1D3G; 15102S; Cell Signaling Technology)

#### Secondary antibodies

Alexa Fluor 488-conjugated goat anti–mouse (1:1,000; A-11029; Thermo Fisher Scientific)

Alexa Fluor 488-conjugated goat anti–rabbit (1:1,000; A-11034; Thermo Fisher Scientific)

Alexa Fluor 647-conjugated goat anti–mouse (1:1,000; A-21236; Thermo Fisher Scientific)

Alexa Fluor 647-conjugated donkey anti–rabbit (1:1,000; A-31573; Thermo Fisher Scientific)

### Live cell imaging

Cells were plated in 8-well glass bottom μ-slides (ibidi) at least 24 hr prior to imaging. For siRNA experiments, cells were transfected in the chamber slide and hydroxyurea was added at least 4 hr after last transfection, and most often 24 hr after. Live cell imaging was performed using a 20×/0.70 Plan Apo Leica objective or a 40x/1.30 Plan Apo Leica objective (where noted) on an automated Leica DMi8 microscope outfitted with an Andor CSU spinning disk unit equipped with Borealis illumination, an ASI automated stage with Piezo Z-axis top plate, and an Okolab controlled environment chamber (humidified at 37°C with 10% CO2). Long term automated imaging was driven by MetaMorph software (version 7.10.0.119). Images were captured with an Andor iXon Ultra 888 EMCCD camera. For most experiments, cells were imaged every 3 min for 24 hr. For analysis of GFP-BAF and RFP-NLS kinetics and the duration of very short nucleus ruptures, images were acquired every 30 sec for 4 hr (30 sec pass time). For micronucleus experiments cells were imaged every 3 min for 36 hr. These exceptions are noted in the figure legends. Post-processing of image stacks was performed in FIJI. Where noted, RFP-NLS or GFP-NLS signal was gamma adjusted to enhance visibility of cytoplasmic FP-NLS signal.

### Laser-induced rupture

Laser induced rupture was performed with a Leica HCX Plan Apo 63x/1.40 Oil CS2 objective on an automated Leica TCS SP8 microscope equipped with adaptive focus control. Cells were imaged in phenol red free DMEM (GIBCO) with 25mM HEPES and 10% FBS (GIBCO). An LCI stage top environment control chamber maintained cells at 37°C, 5% CO2, with humidity. Laser ablation was controlled by MetaMorph (Molecular Devices) using an Andor Micro-Point laser tuned to 435 nm and images were acquired using LAS X software (Leica). RFP-NLS was imaged with a white light laser tuned to 561 nm and a pinhole set to 6.0 AU to enhance signal detection. Nuclear rupture was induced by targeting 6 laser pulses (approximate duration of 30 nsec) to a single spot (approximate ablation diameter of 220 nm) on the nuclear rim. Targeted cells were selected based on average RFP-NLS expression levels and normal nucleus morphology and non-herniated nucleus poles were targeted for ablation. Hernia were avoided to reduce variability in NE composition at ablation sites Nuclei were imaged every 3 min for 5 frames prior to laser targeting and 12 hr after. Post-targeting imaging was initiated within 3 min after the laser pulses. Laser-induced rupturing was optimized on cells expression GFP-BAF. Only spontaneous ruptures observed during imaging of laser-induced ruptures were included in the associated analyses.

### Image analysis

All image analysis was performed in FIJI and graphing performed in Prism 8 (GraphPad). For RFP-NLS intensity analysis, images were background subtracted using the rolling ball method (r = 150px, sliding parabola), and the mean intensity of all nuclei in the field was analyzed for each frame by applying a mask based single global threshold optimized for each image sequence. Background subtracted image sequences were then cropped to individual nuclei and the RFP-NLS mean intensity for that nucleus was analyzed by selecting and analyzing a large ROI in the middle of the nucleus in each frame. These values were then normalized to the corresponding field RFP-NLS mean intensity for each frame. For individual ruptures, these values were then normalized the mean RFP-NLS mean intensity of the 5 frames prior to nucleus rupture and the first drop in RFP-NLS intensity (rupture start) set to t = 0. Rupture extent was quantified on traces where all values were normalized to the mean of the 5 RFP-NLS intensity measurements prior to rupture (t = −15 min to t = −3 min) and calculated as initial RFP-NLS intensity (mean) – RFP-NLS intensity (MIN). RFP-NLS intensity loss was quantified as (time of first consistent increase in RFP-NLS intensity post rupture) + 3 min and RFP-NLS intensity regain was quantified as (time when intensity meets or exceeds the mean of the 5 RFP-NLS intensity measurements after rupture ends) - (time of first consistent increase in RFP-NLS intensity post rupture).

For GFP-BAF integrated density analysis, image stacks were cropped to rupturing nuclei or nuclei plus rupturing micronuclei, then a Bernsen local thresholding algorithm was used to identify and segment GFP foci. Foci integrated density was measured for as long as the object could be identified from background or from nearby ruptures. Where indicated, GFP integrated density values were normalized to the maximum value. For figure 1F, GFP-BAF mean intensity measurements were acquired for 5 frames prior to rupture using the area of the first observed foci as the ROI for the primary nucleus and the entire RFP-NLS signal as the ROI for the micronucleus. The average (mean) GFP-BAF of these 5 frames was used as the “pre-rupture” intensity. The “post rupture” intensity was the last measurement in which the foci was distinguished from the background by thresholding. For comparison of RFP-NLS extent and GFP-BAF recruitment, the maximum GFP-BAF integrated density measurement for each rupture was used.

Rupture duration was quantified by counting the number of frames where RFP-NLS was visible in the cytoplasm and multiplying by 3 min or 0.5 min based on the image acquisition frequency (3 min vs. 30 sec). To be analyzed, nuclei had to satisfy the following criteria: (1) normal nucleus morphology with the exception of chromatin herniations or membrane blebs, (2) RFP-NLS is completely reimported into the nucleus within the imaging window, (3) nucleus is present in frame for at least 75% of imaging duration, and (4) the cell did not undergo apoptosis. The proportions of nuclei that apoptosed and failed to repair were quantified for each experiment, and no significant differences between conditions was observed (data not shown). For experiments with GFP-BAF mutants, only GFP positive cells were included in the analysis. Cells were selected at random from multiple fields of view for each experiment. To eliminate bias, either a subset (>25%) or the entire dataset was blinded prior to analysis. For analysis of protein recruitment to rupture sites in fixed images, the presence of fluorescent protein puncta with a fluorescent signal greater than the surrounding NE was assessed in cells with cytoplasmic RFP-NLS.

### Statistical analysis

All statistical tests were executed in Prism. For comparison of rupture duration total populations, a non-parametric Kolmogorov-Smirnov (K-S) test was used to assess significant differences in the median or distribution of rupture durations between groups. The Kruskai-Wallis (K-W) test was applied as a one-way Anova family test for comparison between 3 or more conditions followed by Dunn’s multiple comparison correction. For comparison of rupture duration histograms, a family chi-square test was performed on pooled data from multiple experiments to evaluate the propriety of individual value testing using the null hypothesis that the populations are the same. For assessing differences between conditions for an individual rupture duration, a Fisher’s exact test was performed after pooling numbers from all replicates comparing the number of ruptures in the category versus all other ruptures. Pooled values were required for statistical testing to ensure that no “0” or other very low values were present. Graphs of nominal data depict proportions derived from individual experimental replicates (dots) and proportions after pooling all replicates (bars) to display variability between experimental replicates. Additional statistical tests used in the manuscript are described in the figure legends. For all tests, a p value less than or equal to 0.05 was considered significant.

## Abbreviations

BAF: barrier-to-autointegration factor
K-S: Kolmogorov-Smirnov test
K-W: Kruskal-Wallis test
LEM: Lap2, emerin, Man1
LEMD2: Lem domain-containing protein 2
MN: micronucleus
NE: nuclear envelope
aaRS: aminoacyl-tRNA synthetase
NETs: nuclear envelope transmembrane proteins
NLS: nuclear localization signal
PN: primary nucleus

## Acknowledgements

We would like to thank Dan Gerlich’s lab at the Institute of Molecular Biotechnology of the Austrian Academy of Sciences for the BAF constructs. E.M.H is supported by the National Institutes of Health grant R35GM124766-02. This research was supported by the Cellular Imaging and Flow Cytometry Shared Resources of the Fred Hutch/University of Washington Cancer Consortium (P30 CA015704). A.M.Y. is supported by the National Institutes of Health postdoctoral fellowship F32CA232764-01.

---

**Figure S1.**
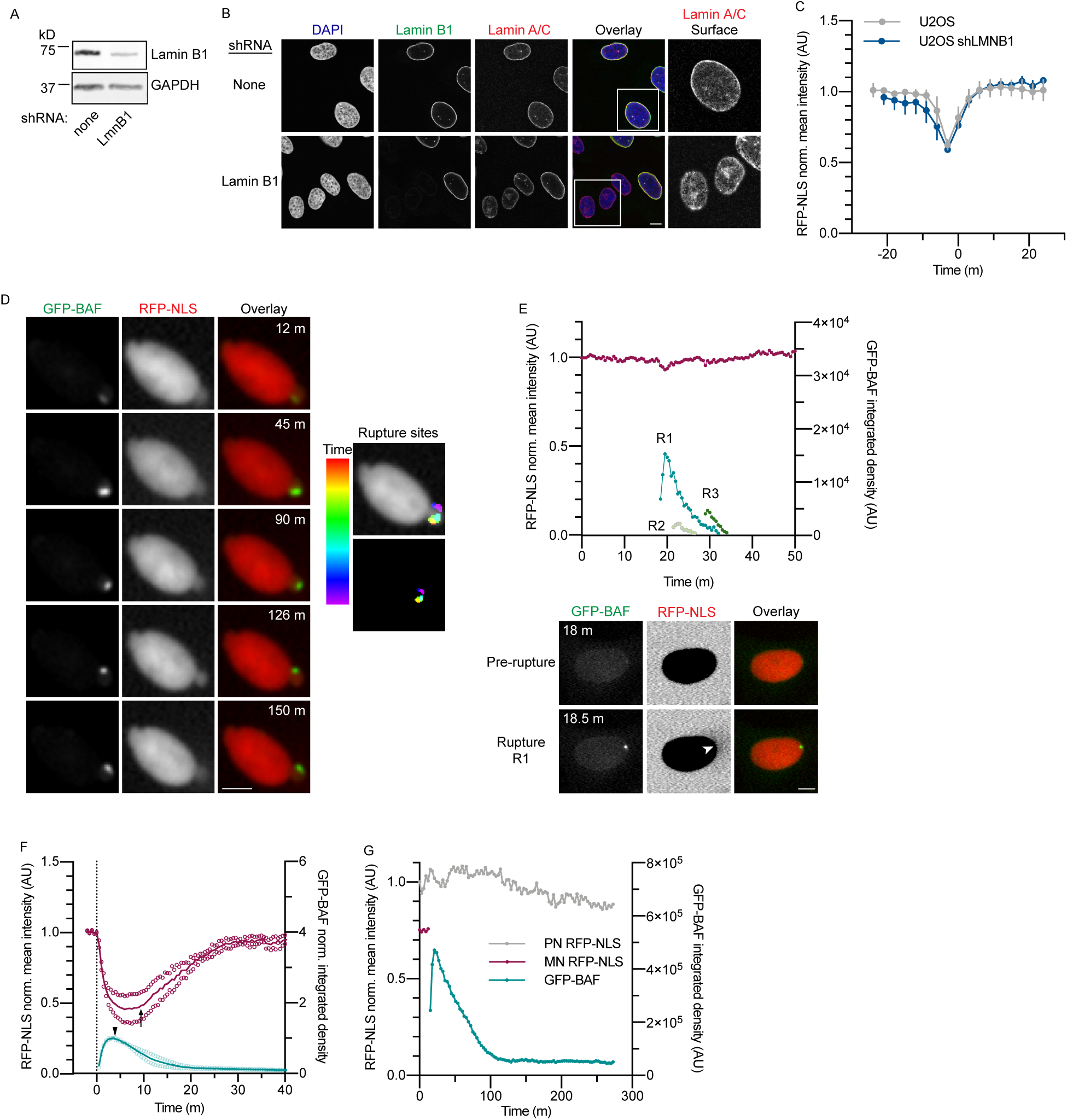
Characterization of U2OS shLMNB1 cell line, BAF recruitment to clustered ruptures, and kinetics of GFP-BAF loss from rupture sites. (A) Representative immunoblot of lamin B1 protein levels in normal versus LMNB1 shRNA expressing U2OS cells. Mean lamin B1 expression in shLMNB1 cells relative to control was 0.25, n = 3 blots. (B) Representative images of U2OS and U2OS shLMNB1 cells labeled with antibodies to lamin B1 and lamin A/C. Images are a single slice from the center or bottom (“surface”) of the nucleus. Boxes indicate nuclei enlarged in surface slices, which show an increase in nuclear lamina gaps in cells depleted of lamin B1. (C) Quantification of RFP-NLS mean intensity during nucleus ruptures (extent rupture = 0.3 to 0.5) in U2OS and U2OS shLMNB1 cells (n values: U2OS, 15; U2OS shLMNB1, 6 ruptures). t_0_ = start of repair (first value of consistent RFP-NLS intensity increase). Mean and 95% CI plotted. (D) Still images of multiple ruptures on the same herniation in a U2OS GFP-BAF RFP-NLS shLMNB1 repair cell, 30 sec pass time. Each row depicts a separate rupture event and GFP-BAF accumulation. Right: GFP-BAF images from left panels are overlaid and color coded for time of rupture. (E) Top: RFP-NLS mean intensity and GFP-BAF integrated density traces for three ruptures in a single nucleus (R1-3) occurring within 11 min of each other, 30 sec pass time. Bottom: still images of rupture 1, with RFP-NLS intensity inverted and oversaturated to show RFP-NLS in cytoplasm concurrent with GFP-BAF accumulation. (F) RFP-NLS mean intensity and GFP-BAF integrated density traces aligned and normalized as in figure 1C, for ruptures where duration > 5 min, 30 sec pass time. Mean (thick line) and replicates (open circles) are plotted. Arrow = time of repair start, arrowhead = time of first GFP-BAF decrease. n = 2 ruptures. (G) RFP-NLS mean intensity and GFP-BAF integrated density traces from the primary nucleus (PN) and micronucleus (MN) of the cell in figure 1F. RFP-NLS is absent from the MN after membrane rupture. Scale bars = 10 μm. AU = arbitrary units.

**Figure S2.**
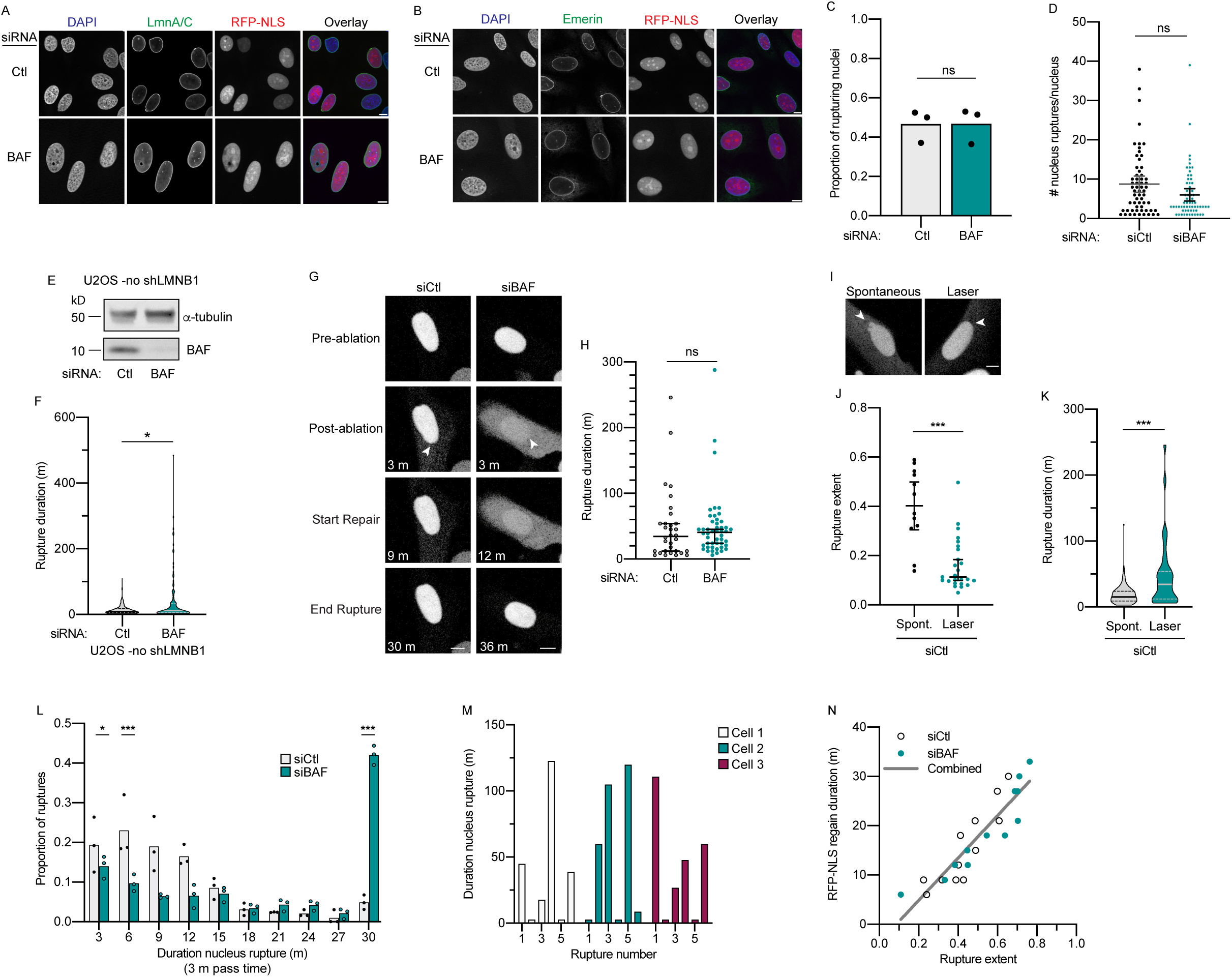
Additional characterization of BAF depletion phenotypes. (A-B) Representative images of U2OS RFP-NLS shLMNB1 cells 48 hr after transfection with siControl (Ctl) or siBAF and labeled with antibodies to lamin A/C and emerin. Scale bars = 10 μm. (C) Quantification of the proportion of nuclei that ruptured at least once during 24 hr of imaging after transfection with siCtl or siBAF (n values: siCtl, 597; siBAF, 616 ruptures from 3 experiments. ns = p > 0.05, Fisher’s exact test. (D) Quantification of mean ruptures/nucleus (n values: siCtl, 61; siBAF, 64 nuclei from 5 and 4 experiments, respectively). (E) Representative immunoblot of BAF protein levels in U2OS cells 48 hr after siRNA transfection. (F) Quantification of nucleus rupture durations after transfection with siCtl or siBAF (n values: siCtl, 165; siBAF, 124 ruptures from 3 experiments). *p<0.05, K-S test. (G) Still images of RFP-NLS from U2OS RFP-NLS shLMNB1 cells transfected with indicated siRNAs after laser-induced rupture. Start repair = RFP-NLS regain start in figure 2A. (H) Quantification of nucleus rupture durations in cells transfected with indicated siRNAs after laser induced rupture (n values: siCtl, 32; siBAF, 48 ruptures over 3 experiments). ns = p>0.05, K-S test. (I) Still images of RFP-NLS from cells transfected with siCtl showing maximum extent rupture during either spontaneous or laser-induced nucleus ruptures. Arrowheads in J-K indicate rupture sites and scale bars = 10 µm. (J) Quantification of nucleus rupture extent during either spontaneous (Spont.) or laser-induced nucleus ruptures (n values: spontaneous, 12; laser-induced, 25). ***p<0.001, Mann-Whitney test. (K) Quantification of nucleus rupture duration of spontaneous or laser-induced ruptures in cells transfected with siCtl (n values: spontaneous, 126; laser-induced, 32). ***p<0.001, K-S test. (L) Histogram of proportion of ruptures shown in Figure 2C with indicated nucleus rupture durations. ***p<0.001, *p<0.05, Fisher’s exact test. (M) Nucleus rupture duration, in order of rupture, for 3 representative cells undergoing at least 6 ruptures during imaging. (N) RFP-NLS intensity regain duration compared to rupture extent from siCtl and siBAF cells, both quantified from RFP-NLS intensity trace analysis (n values: siCtl, 10; siBAF, 9 ruptures). Best fit line for the combined data sets is shown.

**Figure S3.**
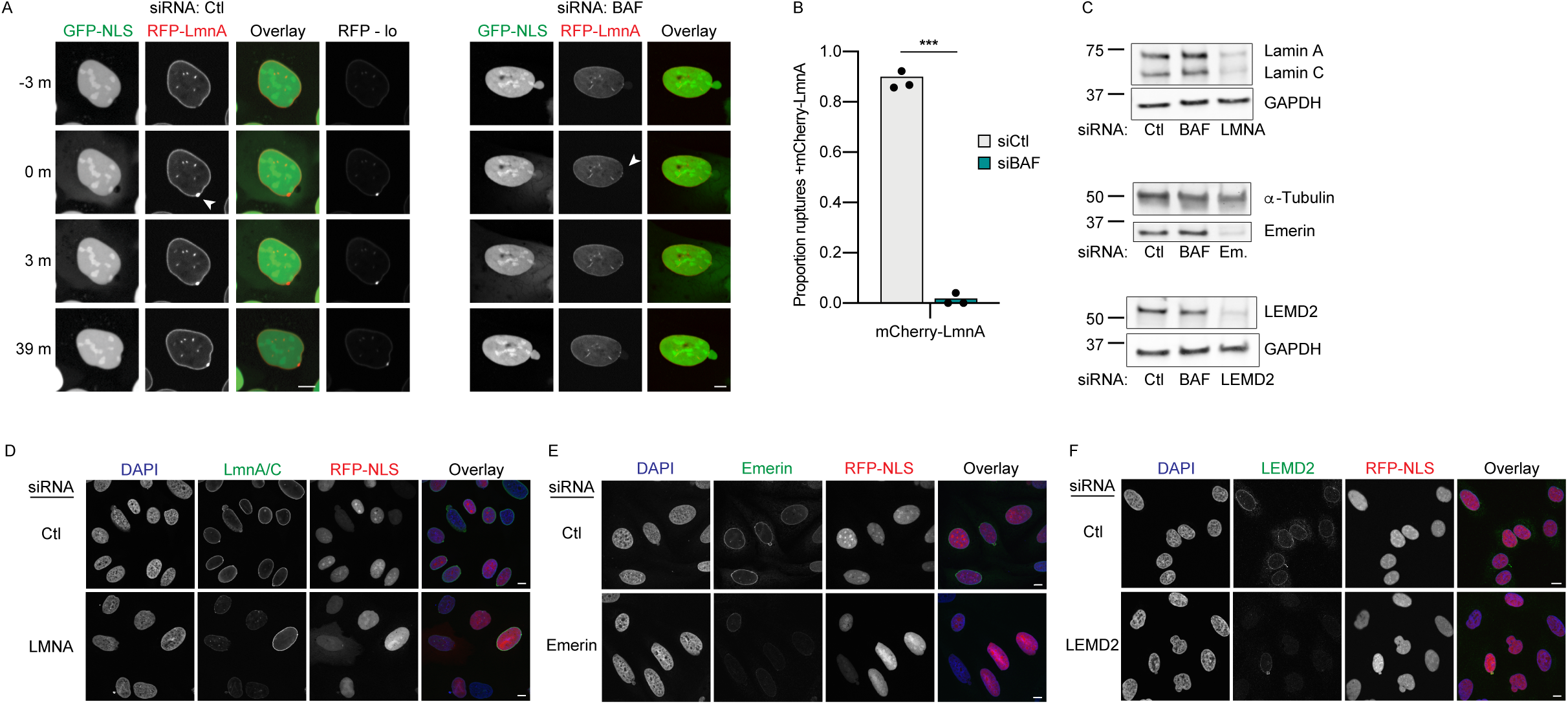
BAF is required for lamin A recruitment to rupture sites and controls for NE protein knockdowns. (A) Still images of U2OS cells stably expressing mCherry-Lamin A (RFP-LmnA), 3xGFP-NLS, and shLMNB1 48 hr after transfection with siControl (siCtl) or siBAF. Cells were imaged at 40x every 3 min for 24 hr. RFP-lo are unsaturated versions of the indicated RFP-LmnA images. Arrowheads mark membrane rupture site. (B) Quantification of rupture sites with mCherry-LmnA protein accumulation. N values: siCtl, 63; siBAF, 53 ruptures from 3 experiments. ***p<0.001, Fisher’s exact test. (C) Representative immunoblot of NE protein levels after 48 hr siRNA treatment. Mean values of targeted protein expression compared to control are siLMNA: 0.24 (n = 2 blots), siEmerin: 0.18 (n = 4), siLEMD2: 0.49 (n = 3). (D-F) Representative images of protein depletions in cells transfected with indicated siRNAs and labeled with antibodies against the targeted protein. All FP-NLS images are gamma adjusted. All scale bars = 10 μm.

**Movie S1.** Time lapses of nuclear membrane rupture and repair in cells expressing GFP-BAF or after BAF depletion, related to Fig. 1A and D, and Fig 2C. First Video: GFP-BAF localization in U2OS RFP-NLS shLMNB1 cells undergoing primary nucleus (PN) and micronucleus (MN) membrane ruptures. RFP-NLS nucleus intensity decreases upon rupture (red) concurrent with GFP-BAF (green) accumulation at rupture site. (Left) Cells were arrested in S phase and imaged every 30 sec. Rupture corresponds to R1 in figure1, A and B. (Right) Cells were treated with reversine then imaged every 3 min. Both movies are played at 1 sec = 7.5 min real time. Second Video: U2OS RFP-NLS shLMNB1 cells transfected with control or anti-BAF siRNAs, arrested in S phase, and RFP-NLS imaged every 3 min. In movie, 1 sec = 45 min. All videos acquired at 20X. Time stamps are hh:mm:ss. Scale bar is 10 μm.

## Notes

### Competing Interest Statement

The authors have declared no competing interest.

